# Peptide-Induced Formation of Extracellular Vesicles that are Distinct From Endogenous *E. coli* OMVs, and Provide an Enhanced Platform for Protein Production and Purification

**DOI:** 10.64898/2026.03.12.711285

**Authors:** Bree R. Streather, Tara A. Eastwood, Karen Baker, Mingzhi Liang, Alexandra E. Bailie, Tijn T. van der Veiden, Lars J. C. Jeuken, Stanley W. Botchway, Lin Wang, Daniel P. Mulvihill

## Abstract

Bacterial outer membrane vesicles (OMVs), are nano-sized, spherical structures released by Gram-negative bacteria that play diverse roles in bacterial physiology, including communication, nutrient acquisition, and host interactions. These vesicles bud from the bacterial outer membrane and contain lipopolysaccharides, periplasmic proteins, nucleotides, and other biomolecules. The Vesicle Nucleating peptide (VNp) is a short peptide tag that, when fused to the amino terminus of a protein of interest, promotes the formation of bespoke recombinant extracellular vesicles in *Escherichia coli*, enabling efficient production and simplified purification of recombinant proteins. Here, we characterise VNp-induced vesicles and compare their composition and organisation with naturally produced *E. coli* OMVs. While both vesicle types possess a single outer membrane-derived lipid bilayer, recombinant protein is highly enriched within the VNp vesicles compared to endogenous OMVs. VNp-fusions and periplasm-targeted recombinant proteins localize to distinct vesicle populations, with VNp-fusions showing markedly higher intra-vesicular concentrations and vesicular purity, compared to the OMV targeted protein. OmpX co-expression further enriched the VNp-fusion content of vesicles, further enhancing yield. The VNp-vesicle lumen is an oxidizing environment, thus supports formation of inter- and intra-molecular disulfide bonds within encapsulated proteins. Overall, VNp-induced vesicles represent a distinct class of recombinant extracellular vesicles that offer a simple and efficient route for producing and purifying concentrated, correctly folded recombinant proteins, expanding the utility of bacterial vesicle systems for biotechnological applications.

**Significance Statement:** Bacterial extracellular vesicles are recognized as versatile tools for biotechnology yet engineering bacterial vesicle production in a controlled and efficient manner remains challenging. Here we describe how a short Vesicle Nucleating Peptide (VNp) tag, fused to a protein of interest, that can be used to program *Escherichia coli* to produce recombinant extracellular vesicles that are compositionally and structurally distinct from natural bacterial outer membrane vesicles (OMVs). VNp-induced vesicles are more homogeneous, and more highly enriched in target fusion proteins, providing a simple and efficient route for protein production and purification. The oxidizing lumen of these vesicles supports disulfide bond formation, and rapid compartmentalisation enables expression of otherwise challenging or toxic proteins. This work characterises a distinct class of recombinant bacterial vesicles and establishes a practical platform for producing correctly folded, concentrated, partially purified proteins in a self-packaged form, expanding the potential applications of bacterial extracellular vesicles in biotechnology and synthetic biology.

## Introduction

All Gram-negative bacteria are capable of releasing cellular material into spherical membrane-bound structures known as outer membrane vesicles (OMVs) (1). They form when a region of the bacterial outer membrane (OM) swells outwards, forming a protuberance that pinches off, without a loss of membrane integrity, and are subsequently released into the extracellular medium. Since their discovery in 1965 (2), various functions have been hypothesised including cell-cell communication; virulence factor delivery to host cells; horizontal gene transfer and nutrient breakdown. OMV production is actively regulated by the bacteria in response to cellular stress, nutrient status, and interaction with host environments. As well as OM lipids and associated lipopolysaccharides (LPS), OMVs also contain periplasmic proteins, nucleic acids, signalling toxins and, rather than simply being filled with indiscriminate periplasmic content, are enriched for certain proteins (3).

It is evident that OMVs play a key role in pathogenicity, primarily through the targeted release of virulence factors, delivered to host cells by endocytosis (4, 5). Furthermore, with a membrane composition similar to that of the OM, they can help bacteria to evade the host immune system by acting as a decoy to immune cells. The main immuno-stimulatory component of the OM is LPS which makes up the majority of the outer leaflet making it highly accessible for detection. This molecule consists of a lipid A core, the outer core non-repeating oligosaccharide and the O-antigen, another polysaccharide (6). The structure of lipid A and the O-antigen varies between bacterial strains and species but the core oligosaccharide tends to be well conserved within a genus or family showing its importance in membrane integrity. The lipid A moiety can be detected at picomolar concentrations by the toll-like-receptor 4 (TLR4) in the host innate immune system causing the release of inflammatory cytokines and activation of many downstream processes, potentially leading to endotoxic shock which can be fatal. The O-antigen triggers the complement cascade but a longer O-antigen can confer resistance to the effects of this (7). LPS has been identified in OMVs produced from *Pseudomonas aeruginosa* (8), *Porphyromonas gingivalis* (9) and *Escherichia coli* (10). Alongside the detection of LPS, the phospholipid composition of OMVs has also been analysed. In those produced from *E. coli*, this is reported to be comparable to that of the OM with ∼80% phosphatidylethanolamine (PE), 15% phosphatidylglycerol (PG) and 5 %cardiolipin also present (4, 11). Compared to the phospholipid composition of whole *E. coli* cells (75% PE, 20% PG and 5% CL (12)), there is a slightly elevated proportion of PE.

OMV production also has physiological relevance for non-pathogenic bacteria, with a role in nutrient acquisition (13), achieved by preferentially packaging certain enzymes into OMVs (14). Hypervesiculation can occur as a result of stress, particularly in the case of accumulation of peptidoglycan fragments in the periplasm (15), LPS in the OM (16) or toxic/misfolded proteins (17). Some knockout strains have a hypervesiculation phenotype, the discovery of which has led to the development of an *E. coli* strain, based on the BL21(DE3) strain used for recombinant protein production, which is able to produce up to three-fold more OMVs. This strain, BL21(DE3)Δ60, has been shown to be capable of targeting recombinant proteins to the vesicles (18).

This example reflects a broader trend in biotechnology of exploiting bacterial OMVs, particularly in vaccine development, due to their ability to elicit an immune response and their potential as targeted antigen delivery systems. For this, translocation of the recombinant protein into the periplasm is required, through the use of an appropriate signal peptide. Proteins that must fold in a reducing environment are transported via the Tat pathway, fully folded, whereas those which require disulfide bond formation, an oxidising process, are transported in an unfolded state through the Sec pathway (19). This dependence on periplasmic disulfide bond formation, catalysed by the DsbA/DsbB system (20), represents a fundamental limitation of recombinant protein production in *E. coli*.

This, and other limitations including issues with low solubility and yield, can be overcome by using the Vesicle Nucleating peptide (VNp) technology (21). This innovative method uses a short peptide tag, derived from human synucleins to target recombinant proteins into membrane-bound vesicles. These vesicles can be either extracellular or intracellular. VNp fusions significantly increase the yield of a protein of interest (POI) with several VNp variants having been produced to maximise POI yield. VNp2, VNp6 and VNp15 are currently the most commonly used due to their protein-packaging efficiency (22). This technology also allows for the production of otherwise toxic or insoluble proteins, including disulfide bond-containing antibodies (23), through compartmentalisation. The extracellular vesicles provide a stable storage environment and simplify downstream processing with the POI being the most abundant protein contained within them. Bacterial cells can be easily removed from the culture by centrifugation, leaving vesicles in the supernatant and reducing the number of contaminants. Purified recombinant proteins from VNp vesicles are properly folded and functional. Its many advantages make VNp attractive for numerous applications across biotechnology.

VNp has proven successful with a wide range of POIs but little is known about the biophysical properties of VNp vesicles, their relationship to innate OMVs or their mechanism of formation. Currently, it is predicted that in the formation of extracellular vesicles, VNp interacts with the bacterial membrane to induce outward membrane curvature which continues until the membrane fuses back on itself, the vesicle buds off and is released into the media. It is unclear if it first interacts with the inner membrane (IM) causing subsequent bulging of both membranes and resulting in vesicles with two membrane layers or whether the VNp fusion is translocated to the periplasm before inducing outward curvature of the OM only, like endogenous OMVs (eOMVs). The determining factor for intracellular or extracellular vesicle formation is also unknown.

Here we describe a detailed biophysical characterisation of VNp-fusion induced vesicles and compare them with natural bacterial OMVs. We find that while they share some similar lipid composition and both contain oxidising environments, there are several key differences. The intra-vesicular concentration of VNp-fusion is significantly higher than that observed for an equivalent periplasmic targeted protein. The higher intra-vesicle concentration in part explains why VNp-fusion induced vesicles provide higher purity and protein yields compared to OMVs, which not only simplifies downstream purification, but results in significantly higher yields of the target protein. These properties make them extremely attractive vehicles for the simple production and isolation of a wider range of recombinant protein than is normally allowed with the *E. coli* host system and have the potential for a wide range of biotechnology applications.

## Materials and Methods

### Bacterial cell Culture and protein induction

BL21 Star (DE3) *E. coli* cells (F^-^*omp*T *hsd*S_B_ (r_B_^-^, m_B_^-^) *gal dcm rne131* (DE3)) were used for all experiments. *E. coli* cells were cultured using LB (10 g Tryptone; 10 g NaCl; 5 g Yeast Extract (per litre)) and TB (12 g Tryptone; 24 g Yeast Extract; 4 ml 10% glycerol; 17 mM KH_2_PO_4_ 72 mM K_2_HPO_4_ (per litre)) media each supplemented with kanamycin (50 µg/ml final conc.). 5 ml LB starters from fresh bacterial transformations were cultured at 37°C to saturation and used to inoculate 25 - 500 ml volume TB media flask cultures that were incubated overnight at 30°C with 200 rpm orbital shaking (2.5 cm throw). Oxygenation of cultures was maximised, to promote vesicle production, by maintaining a large liquid surface area during growth (e.g. 25 ml media in 500 ml conical flask or 500ml / 1 L of media in a 5 L conical flask). Recombinant protein expression from the T7 promoter was induced by addition of IPTG to a maximum final concentration of 20 µg/ml (84 µM) once the culture had reached an OD_600_ of 0.8 - 1.0. It should be noted that use of higher IPTG concentrations results in cell lysis. Growth curves were generated from 96 well plate cultures, prepared from late log-phase cultures, diluted into fresh media to an OD_600_ of 0.1at the start of the growth analysis experiment. Fluorescence signal from overnight plate cultures were obtained using a BMG Labtech Clariostar plus shaking UV & fluorescence plate reader. Plates were incubated at 37 °C using double-orbital shaking, and 589 nm absorbance values were taken every 15 minutes for the duration of the experiment. Fluorescence curves were generated from averages of 4 individual biological repeats.

### Molecular Biology

All tags and gene sequences were codon optimised for expression in *E. coli,* and synthesised DNA sequences (*Thermofisher*) were cloned into pRSFDUET-1 plasmid (*Novagen*) using appropriate restriction enzymes. Bacterial expression plasmids and sequences described in this study (pRSFDUET-VNp6-mCherry2 (v1265); pRSFDUET-OmpX-mNeongreen (v1800); pRSFDUET-VNp6-mCherry2_OmpX-mNeongreen (v1640); pRSFDUET-ssDsbA-mCherry2 (v1596); pRSFDUET-ssDsbA-mCherry2_OmpX-mNeongreen (v1650); pRSFDUET-ssDsbA-mCherry2 VNp6-mNeongreen (v1670); pRSFDUET-VNp6-mCherry2-StefinA (v1746); pRSFDUET-VNp6-mCherry2-Uricase (v1745); pRSFDUET-VNp6-mNeongreen (v1138); pRSFDUET-ssDsbA-mNeongreen(v1595) are available from Addgene upon publication (https://www.addgene.org/Dan_Mulvihill/). Note: Stable recombinant extracellular vesicle production requires use of plasmids with antibiotic selections that do not target the cell wall.

### Recombinant vesicle isolation

To isolate vesicles from VNp-fusion and natural OMVs from cultures, *E. coli* cells were initially pelleted from cultures by centrifugation at 3,000 xg for 30 mins. The subsequent vesicle containing media was passed through a 0.45 µm polyethersulfone (PES) filter and either analysed directly, stored at 4 °C or concentrated using a 20 ml 50 kDa cutoff PES centrifugal concentrator (Fisher Scientific) by centrifugation at 500 xg to reduce volume from 25 ml to 1 ml.

### Protein concentration determination

Fluorescence scans were used to determine concentration of mCherry2 fusion proteins in vesicle containing media and soluble protein extracts. Absorbance was measured at 589 nm using a Varian Cary 50 Bio UV-Vis spectrophotometer, with measurements from an equivalent empty vector culture used for baseline correction. Concentration was determined using an extinction coefficient of 79,400 M^-1^cm^-1^.

### Preparation and analysis of phospholipids

Equivalent cell mass samples were isolated by centrifugation from timepoints during overnight *E. coli* cultures, chilled on ice and then rapidly frozen. Vesicles were isolated and concentrated as described above. Lipids were isolated by adding 300 µl of methyl-tert-butyl-ether (MTBE): MeOH (10:3, v/v) per 50 mg cell pellet, mixed rapidly by shaking and vortex, prior to incubation continuous shaking at RT for 1 hour. dH_2_O was added to a final volumetric ratio of 10:3:2.5 (MTBE: MeOH: dH_2_O), then vortexed and incubated with continuous shaking at RT for 1 hour. Samples were subsequently centrifuged at 1000 *xg* for 10 minutes at 4 °C, and the top layer was extracted using a glass Pasteur pipette. The final lipid-containing solvent was evaporated using a rotary evaporator (*Rotavap*) or under a stream of nitrogen, and the resultant dry lipids were stored at −20 °C until required. Prior to Thin Layer Chromatography (TLC), lipid samples were resuspended in either 500 µl (cells extract) or 50 - 100 µl (vesicle extract) of chloroform. The TLC chamber was equilibrated for at least 45 minutes prior to running the TLC by filling to ∼ 1 cm deep with the solvent (65:25:8 v/v chloroform: MeOH: Acetic acid), inserting a piece of filter paper, just smaller than the internal height/width of the tank, prior to replacing the lid, and allowing the TLC chamber to equilibrate. TLC plates (*Merck, Darmstadt, Germany*) were cut to size, a pencil line drawn 1 - 1.5 cm from the bottom and labelled appropriately with samples 1 cm apart (first and last samples at least 0.5 cm from the edge of the plate). Initially, 10 µl of each sample was loaded before determining “equal loading” through densitometry using Image J (24). The plate was placed level on the bottom of the TLC tank, with the solvent below the pencil line and the lid of the tank replaced. Plates were allowed to develop until the solvent front was ∼ 1 cm from the top of the plate, and subsequently allowed to dry fully outside the tank, before staining with aerosolised Molybdenum Blue solution (*Merck, MO, USA*). Glycoprotein staining of SDS-PAGE gels (for detection of lipopolysaccharide (LPS)) was carried out using Pro-Q™ Emerald 300 (25), following the manufacturer’s protocol. NMR analysis of phospholipids purified from VNp-fusion induced vesicles was undertaken as described elsewhere (26).

### Transmitted Electron Microscopy (TEM) thin section analysis of *vesicles*

Vesicles were purified and concentrated as described above. The vesicles (approximately 100µl) were resuspended in 2 ml of 2.5% (w/v) glutaraldehyde in 100 mM sodium cacodylate buffer pH 7.2 (CAB) and fixed for 2 hr at RT with gentle rotating (20 rpm). Vesicles were washed twice into 100 mM CAB by buffer exchange using a 50 kDa cutoff PES centrifugal concentrator (Fisher Scientific) by centrifugation at 500 xg. Vesicles were post-fixed with 1% (w/v) osmium tetroxide in 100 mM CAB for 2 hr and subsequently washed twice with dH2O, by buffer exchange. Vesicles were dehydrated by incubation in an ethanol gradient, 50% EtOH for 10 min, 70% EtOH overnight, and 90% EtOH for 10 min followed by three 10 min washes in 100% dry EtOH. Samples were then washed twice with propylene oxide for 15 min. Vesicles were embedded by re-suspension in 1 ml of a 1:1 mix of propylene oxide and Agar LV Resin and incubated for 30 min with rotation. The vesicles were resuspended in fresh resin and transferred to a 1-mL BEEM embedding capsule, and samples were polymerised for 20 hr at 60 °C. Ultrathin sections were cut using a Leica EM UC7 ultramicrotome equipped with a diamond knife (DiATOME 45°). Sections (70 nm) were collected on uncoated 400-mesh copper grids. Grids were stained by incubation in 4.5% (w/v) uranyl acetate in 1% (v/v) acetic acid for 45 min followed by washing in a stream of dH_2_O. Grids were then stained with Reynolds lead citrate for 7 min followed by washing in a stream of dH_2_O. Electron microscopy was performed using a JEOL-1230 transmission electron microscope operated at an accelerating voltage of 80 kV equipped with a Gatan One View digital camera.

### Cryogenic Electron Microscopy of VNp induced vesicles

Vesicles were purified and concentrated as described above. Quantifoil Lacey carbon 300 mesh Cu grids were glowdischarged for 60 seconds at 15 mA in a PELCO easiGlow device. Vesicles were applied to the grid in a 4 uL sample and blotted with a Vitrobot mark IV, using the following settings 20 blot force, 3 sec blotting, 100% humidity at 4°C. Sample grids were imaged using a Thermofisher Glacios 200 kV microscope equipped with X-FEG electron gun, Selectris energy filter with a 20eV slit width, and Falcon 4i camera. Samples were imaged at 79,000x magnification at a dose of 15 *e*^-^/Å^2^, resulting in images with 0.16 nm pixels.

### Widefield Fluorescence Microscopy

Cells were mounted onto coverslips under < 1 mm thick circular soft 1% agarose pads and attached with appropriate spacers onto glass slides (27), and the remaining gap was flooded (using capillary action) with TB supplemented with kanamycin and 20 µg/ml IPTG. Samples were subsequently visualised on an inverted widefield fluorescence imaging system (28). All live cell imaging was completed within 30 mins of samples being mounted onto coverslips. RADA stain (R&D Systems (Biotechne), MN, USA) was added to samples at a final concentration of 125 µM and incubated for 1 hr at 37 °C before fixation with ethanol.

**Structured Illumination Microscopy (SIM):** was undertaken using a Zeiss Elyra PS 1 microscope with either a 100x NA 1.46 or 63x NA 1.4 oil immersion objective lens (Zeiss α Plan-Apochromat) as described previously (21, 29). Cells and vesicles were mounted as described above, using high precision No.1.5 coverslips (*Zeiss, Jenna, Germany*). A 405 nm laser was used to illuminate FROG/B, while 488 nm and 561 nm laser were used to illuminate mNeongreen and mCherry2 fusions, respectively. The optical filter set consisted of laser blocking filter MBS 405/488/561 as the dichroic mirror, and the dual-band emission filter LBF-488/561. A total of 3 / 5 rotations of the illumination pattern were implemented to obtain two-dimensional information. During time lapse experiments, cells were maintained at 37 °C with constant humidity using an INU incubation system (Tokai Hit, Shizuoka-ken, Japan). Super-resolution SIM image processing was performed using the Zeiss Zen software. Two colour images were aligned using the same software following a calibration using a pre-mounted MultiSpec bead sample (*Zeiss, Jenna, Germany*). Pearsons Correlation coefficient (r) of fluorescence signals was calculated from signal aligned SIM images using the Fiji software (24) coloc2 analysis tool with Costes threshold regression and 10 randomisations (Supplementary Figure 1).

### Fluorescence Lifetime Imaging Microscopy (FLIM)

The one- and two-photon systems used in this work have been previously described (21, 30). Prior to FLIM data acquisition, protein expression levels were verified using confocal microscopy. Here, a Nikon Eclipse C2-Si confocal scan head attached to an inverted Nikon TE2000 or Ti-E microscope (Nikon, Tokyo, Japan) was used. mNeongreen and mCherry2 fluorescent protein (FP) were excited at 491 nm (emission 520/35 nm) and 561 nm (emission 630/50 nm) respectively using an NKT super continuum laser. For both one and two-photon excitation, emission was collected by the same objective through filters (above) and detected with an external hybrid detector module (HPM-100-40, *Becker & Hickl, Germany*), that links a GaAsP photomultiplier tube (PMT) with a time correlated single photon counting (TCSPC) module (SPC-QC-104, *Becker and Hickl, Germany*). Photon counts of at least 1000 were used for the multi-exponential analysis. Raw time correlated single photon counting decay curve at each pixel (256 x 256 or higher) of the images were analysed using SPCImage software v.8.8 (*Becker and Hickl, GmbH*); a single exponential fit model with a laser repetition time value of 25 ns was used for the decay curve fitting (Supplementary Figure 2). *Vesicular concentrations* of mCherry fusions were determined as described previously (31, 32) using a one photon excitation FLIM system, equipped with a SuperK EXTREME NKT-SC 470-2000 nm supercontinuum laser (NKT Photonics) which generates at variable repetition rate with 70 ps pulse width. The desired wavelengths were selected using a SuperK SELECT multi-line tunable filter (NKT photonics).

Images were collected through a 100X 1.49 NA TIRFM oil objective (Nikon). Laser power was set at 20%, and images were captured using a large pinhole, 20 µm field zoom, 5 µsec pixel dwell time, over a 512 x 512 pixel area. Image data was analysed with binning set at 3. A photon# - FP concentration calibration equation was generated from photon measurements averaged from 5 separate FLIM acquisitions of samples of known concentrations of FP dissolved in PBS. To ensure consistency between calibration solution measurements, data was captured 2 µm above the coverslip. Average photons / pixel gate was measured for ∼ 40 separate vesicles from 3 separate fields of view per sample. Larger objects (typically > 2 x signal from average), likely corresponding to vesicle aggregates, were omitted from the measurements. Average photon values were converted to FP concentration using the previously determined calibration equation. *Anisotropy* values were determined on the above system (using a 60x 1.27 NA water objective lens), as described previously (33). This objective was chosen over the 100x NA 1.49 since the latter is known to introduce large errors for anisotropy measurements. Two-photon excitation anisotropy was employed as the maximum value was 0.57 compared to 0.4 for one-photon excitation. *Viscometry* was determined as described previously (34) on the above multiphoton system using 910 nm excitation. A lifetime - viscosity calibration equation was calculated using equivalent mNeongreen samples dissolved in increasing w/v glycerol solutions (Supplementary Figure 3).

## Results

We have previously described a simple peptide tagging methodology by which the addition of a short VNp sequence to the amino terminal of a protein results in the vesicular export of the resultant recombinant fusion protein from *E. coli* (21). This system has been applied to a wide range of proteins, resulting in high yield production and simplified purification of diverse recombinant proteins (e.g. (23, 35-38)), yet their biophysical organisation and how these vesicles relate to the naturally occurring *E. coli* OMVs remains unknown. To address this, we generated fusions between a bright monomeric fluorescent protein, mCherry2 (mCh2) (39), with either an amino terminal VNp6 tag (MDVFKKGFSIADEGVVGAVEKTDQGVTEAAEKTKEGVM) (21), or a native ssDsbA signalling peptide (MKKIWLALAGLVLAFSASAAQ) that directs co-translational export of proteins through the signal recognition particle (SRP)-dependent translocation pathway (40, 41) into the periplasm from where it is subsequently incorporated into endogenous OMVs (eOMVs). Expression from these VNp6-mCh2 and ssDsbA-mCh2 constructs resulted in the production of mCh2 filled vesicles (Figure 1a & b).

**Figure 1.**
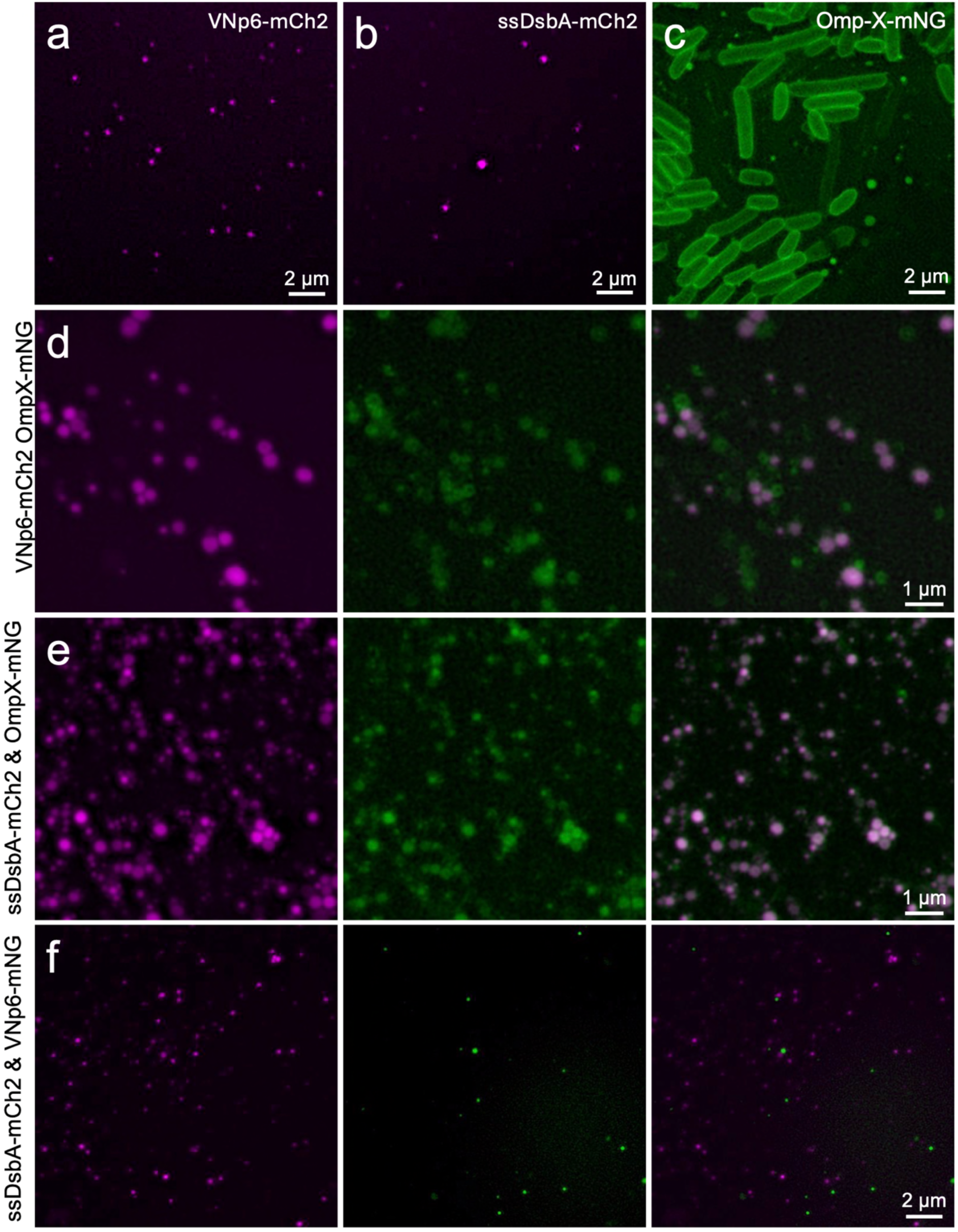
Structured Illumination Microscopy of *E. coli* vesicles. Vesicles isolated from BL21 DE3 cells expressing (a) VNp6-mCh2, (b) ssDsbA-mCh2. (c) A SIM image of a culture of cells expressing fluorescently labelled outer membrane protein, OmpX, highlights the OMV production from *E. coli*. Vesicles isolated from cultures of *E. coli* expressing (d) VNp6-mCh2 and OmpX-mNG, (e) ssDsbA-mCh2 and OmpX-mNG, (f) or ssDsbA-mCh2 VNp6-mNG.

To simultaneously follow vesicle content and outer membrane organisation in cells and vesicles, constructs were generated allowing co-expression of the above proteins with the *E. coli* outer membrane protein OmpX fused to mNeongreen (mNG) (42) at its carboxyl terminus. OmpX is an integral small beta-barrel outer-membrane protein (43), with both amino and carboxyl termini ending in the periplasm. Expression of this construct revealed OmpX-mNG localises to the membrane of bacterial cells and extracellular vesicles within the culture media (Figure 1c). DLS analysis was consistent with the size of vesicle particles in the media. Structured Illumination Microscopy (SIM) timelapse analysis of cells expressing both VNp6-mCh2 and OmpX-mNG revealed mNG labelled vesicles. The VNp vesicles formed rapidly, typically within a minute, and concurrently filled with mCh2 protein (Figure S4).

Vesicles isolated from cells co-expressing OmpX-mNG with either VNp6-mCh2 or ssDsbA-mCh2 possessed OmpX-mNG labelled membranes (Figure 1d-e & S4), with the majority (>95%) of both VNp6-mCh2 and ssDsbA-mCh2 fusion foci seen located within OmpX-mNG labelled vesicles (Figure 1 & S1). While the majority of OmpX-mNG labelled vesicles contained ssDsbA-mCh2 signal (Figure 1e & S1), less than half of OmpX vesicles contained VNp-mCh2 in cultures co-expressing both VNp-mCh2 and OmpX-mNG. (Figure 1d & S1). This indicates the presence of two different population of vesicles, with ssDsbA labelled periplasm targeted to almost all the OmpX-mNG labelled eOMVs, while recombinant VNp-fusion localised to a subpopulation of vesicles, nucleated by the VNp in a mechanism distinct from that resulting in eOMV biogenesis. To test this hypothesis, a construct was generated to allow simultaneous co-expression of ssDsbA-mCh2 (magenta) and VNp-mNG (green) (Figure 1f). In all samples examined, the signals were never observed colocalised within the same vesicle (Figure 1f & S1). These results are consistent with the VNp- and ssDsbA-FP fusions recruiting to separate and distinct, vesicle populations.

Fluorescence lifetime imaging was employed to determine anisotropy values of fusion-proteins within vesicles purified from cultures of *E. coli* expressing VNp-mNG ± OmpX, or ssDsbA-mNG, in order to establish oligomerisation status and dynamics (Table 1). Consistent with previous immuno-EM analysis (21) VNp-fusions were soluble monomers within the vesicle lumen (anisotropy - 0.312 mA ± 0.025) (44), whereas vesicle-associated OmpX-mNG had a lower anisotropy (0.203 mA ± 0.04), consistent with integration within the vesicle membrane (Figure 1c). These anisotropy values are consistent with those previously determined for monomeric and dimeric fluorescently labelled proteins (44). Surprisingly ssDsbA fusions appear as dimers within the vesicle (anisotropy value: 0.229 mA ± 0.081).

**Table 1:** Properties of VNp induced vesicles and ssDsbA labelled OMVs.

| Construct |  | VNp6-fusion | ssDsbA-fusion | VNp6-fusion +<br>OmpX |
| --- | --- | --- | --- | --- |
| Diameter (nm) <sup>†</sup> | : | 163 ± 2.5 | 144 ± 2.6 | 164 ± 1.9 |
| Polydispersity Index value <sup>†</sup> | : | 0.079 | 0.077 | 0.072 |
| Anisotropy of vesicular fusion * | : | 0.312 ± 0.025 | 0.229 ± 0.081 | - |
| Vesicular Viscosity (centiPoise) * | : | 7.677 ± 0.020 | 8.093 ± 0.011 | 6.489 ± 0.024 |
| [ Vesicular mCherry2 ] (μM) ** | : | 7.23 ± 0.83 | 5 ± 0.72 | 9.29 ± 1.304 |
| % total vesicular protein ** | : | 76% | 37% | 85% |
| # mCherry2 molecules / vesicle ** | : | 7.69 | 3.54 | 10.1 |
| mCherry2 in media (μM) ** | : | 9.14 ± 2.17 | 1.39 ± 0.14 | 13.26 ± 0.84 |
| [ Vesicles ] (μM) ** | : | 1.19 | 0.39 | 1.32 |

<sup>†</sup> Determined by Dynamic Light Scattering; \* Determined with mNeongreen fusions; \*\* Determined with mCherry2 fusions.

To provide insight into the origin of the vesicle membranes, we examined the phospholipid composition of the VNp induced vesicles and eOMVs. Samples were taken at timepoints from *E. coli* cells expressing a VNp-fusion protein and vesicles purified from the subsequent culture media. Phospholipids were purified from the samples and analysed by thin layer chromatography (TLC). Molybdenum blue staining of the chromatography plates revealed the presence of phosphatidylethanolamine (PE), phosphatidylglycerol (PG), and cardiolipin (CL) (Figure 2a), the relative abundance of which varied between samples (Figure 2b). As reported previously the relative abundance of CL within *E. coli* increased as the experiment progressed, making up 36% of the membrane lipid in the stationary phase cells (45). In contrast CL made up only 7% the phospholipid composition of VNp induced vesicles. 83% of the vesicle membranes was made up of PE lipid, which is a significantly higher proportion than seen in *E. coli* cells in any growth phase (Figure 2a & b) (4, 11, 12). The relative PG composition of membranes did not vary dramatically between samples. NMR spectra of VNp vesicle phospholipid analysis and comparison with fingerprint spectra of phospholipid standards (26) confirmed the composition and relative abundance of phospholipids within the vesicle membrane (Figure 2d) were consistent with the TLC analysis. When phospholipid composition between natural VNp-fusion induced vesicles, and eOMVs were analysed, the relative lipid composition did not vary significantly between samples (Figure 2c), suggesting a similar cellular origin, and all were consistent with the previously reported composition of the *E. coli* OM (∼80% PE, ∼15% PG, and ∼5% CL) (11).

**Figure 2.**
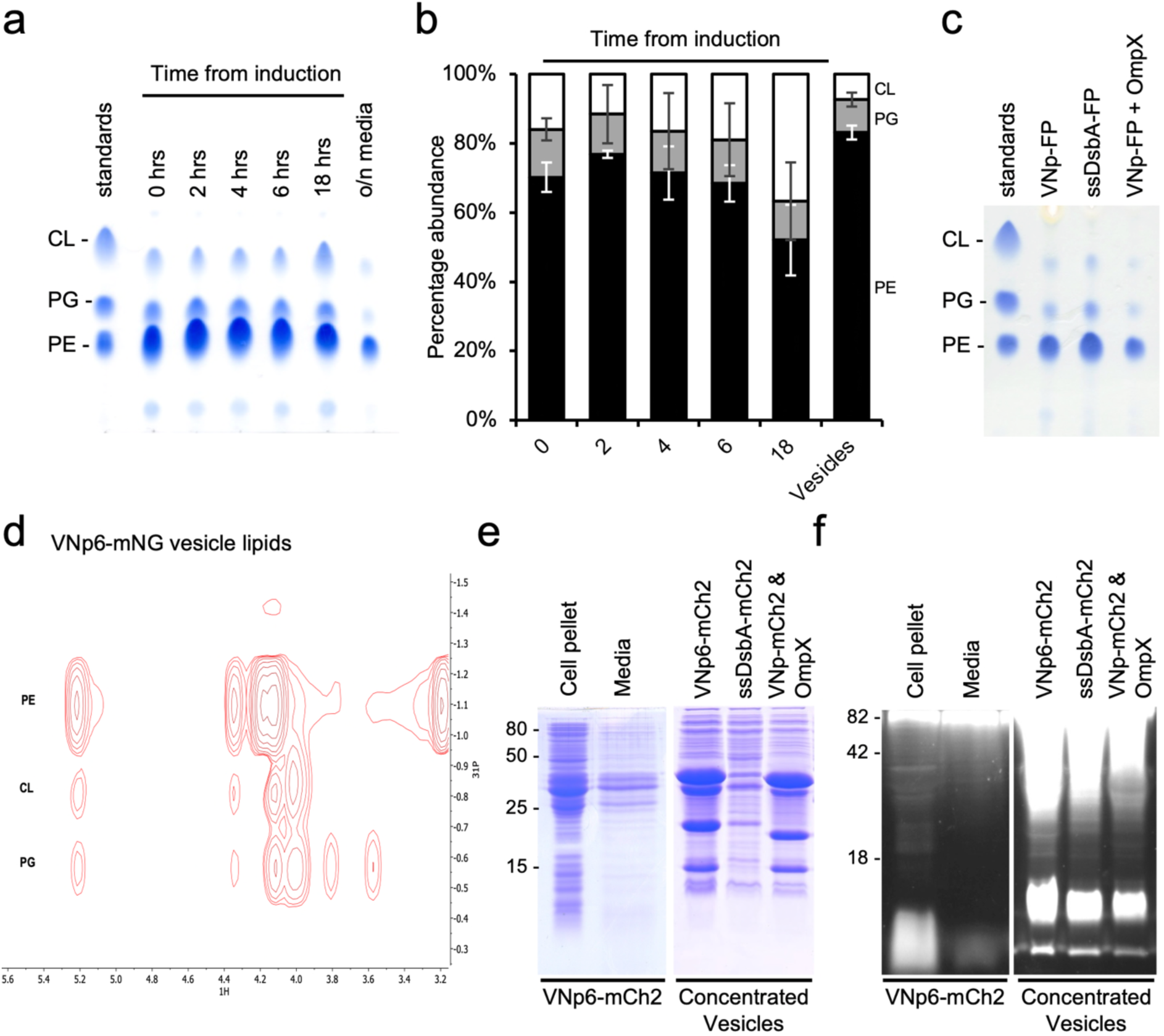
Lipid analysis of VNp-fusion induced vesicles and natural OMVs. (a) Molybdenum blue stained TLC plate with phospholipid extracts from a time course of cells expressing VNp6-mCh2 (time from IPTG induction), and vesicle lipid from the overnight media. (b) Quantification of relative PE (black), PG (grey) and CL (white) phospholipid levels, averaged from 3 biological repeats of experiment show in (a). (c) Molybdenum blue stained TLC plate with phospholipid extracts from vesicles isolated from cultures of *E. coli* expressing VNp6-mCh2, ssDsbA-mCh2, and VNp6-mCh2 OmpX-mNG. (d) 2d NMR spectra of phospholipid extracts from VNp6-mCh2 induced vesicles. Coomassie blue (e) and Pro-Q Emerald 300 (f) stained SDS-PAGE gels of cell pellet and culture media from an overnight culture of *E. coli* expressing VNp6-mCh2, or concentrated vesicles isolated from overnight cultures of bacteria expressing VNp6-mCh2, ssDsbA-mCh2 or VNp6-mCh2 OmpX-mNG.

As an additional analysis of the vesicle membranes, equivalent samples of *E. coli* cells, culture media and vesicle samples were subject to SDS-PAGE. Duplicate gels were stained with either coomassie-G250, to highlight protein composition (Figure 2e), or Pro-Q Emerald 300 (Figure 2f), to confirm the presence of lipopolysaccharide (LPS) and verify the *E. coli* outer-membrane origin of the vesicle membranes. The VNp and ssDsbA fusion bands were present in each sample, but the relative abundance of the two fusions compared to endogenous *E. coli* proteins differed significantly (Figure 2e). In contrast, there was minimal variation in the relative abundance of LPS in each vesicle preparation (Figure 2f), again indicating the membranes from these different vesicles had a common cellular origin.

It is widely reported that the size of eOMVs, as determined by EM and DLS, can vary from 20 - 250 nm in diameter (46-48). Dynamic Light Scattering (DLS) analysis of purified VNp and eOMV vesicles revealed (a) Polydispersity Index values of less than 0.1, indicating the samples were monodisperse, and (b) the VNp-mCh2 vesicles significantly larger than the ssDsbA-mCh2 labelled eOMVs (Table 1). Interestingly co-expression of VNp-mCh2 and OmpX-mNG resulted in no significant difference in vesicle size (163 ± 2.5 nm vs 164 ± 1.9 nm). We went on to examine the membrane organisation and vesicle size distribution of the vesicles using both TEM and cryo-EM imaging (Figure 3). Both TEM images of negatively stained thin sections (Figure 3a & b) and Cryo-EM images of plunge frozen vesicles (Figure 3c) revealed overall membrane organisation was similar between samples, each containing a single lipid bilayer, with a thickness of approximately 5 nm across the constructs examined (Figure 5a), which is consistent with previously determined and published thickness of the *E. coli* outer membrane (49). Interestingly, a significant proportion of vesicles (∼ 10 % overall) contained multiple membrane layers (Figure 5a), which may be a reflection of vesicle formation, or due to an artefact of sample preparation. However, these data indicate that like eOMVs, VNp induced vesicles are encapsulated by membranes originating from the outer *E. coli* membrane.

**Figure 3.**
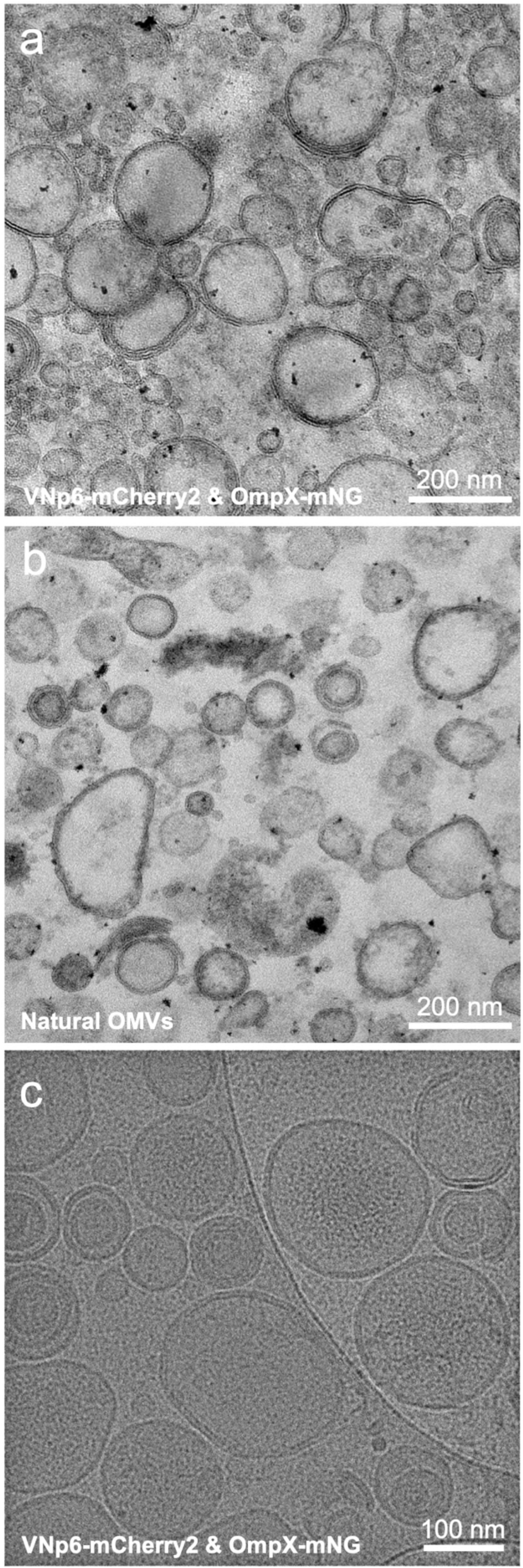
Negative stained thin-section EM analysis of *E. coli* derived vesicles. EM images of negative stained thin sections of vesicles isolated from *E. coli* expressing (a) VNp6-mCh2 and OmpX-mNG or (b) ssDsbA-mCh2. (c) Cryo-EM image of vesicles isolated from *E. coli* expressing VNp6-mCh2 and OmpX-mNG.

To confirm which subcellular region the lumen of VNp induced vesicles originated from, we employed the synthetic biosensor FROG/B (50) to characterise the redox environment within the vesicle lumen. This biosensor has been engineered to emit fluorescence at two wavelengths. While signal in the blue wavelengths is independent of the redox environment and therefore acts as a measure of local protein abundance, intensity in the green increases under oxidising conditions. Vesicles generated from cells expressing VNp-FROG/B contain lumen with a more oxidised environment than the bacterial cytosol (Figure 4a & b). These data are consistent with the presence of disulphide bond containing VNp fusion proteins within both external and internal vesicles (e.g. human growth hormone, Etanercept, monoclonal antibodies, etc (21, 23)). Thus, the measured redox environment within the VNp induced vesicular lumen is consistent with an *E. coli* periplasmic origin. Interestingly, while RADA staining, a tetramethylrhodamine-based fluorescent D-amino acid peptidoglycan dye, highlighted the periplasm of *E. coli* cells, no detectable RADA signal was observed in isolated VNp-mNG induced vesicles (Figure 4c & d), suggesting peptidoglycan is absent from the vesicles, which is consistent with our previous mass-spectrometry based analysis (21).

**Figure 4.**
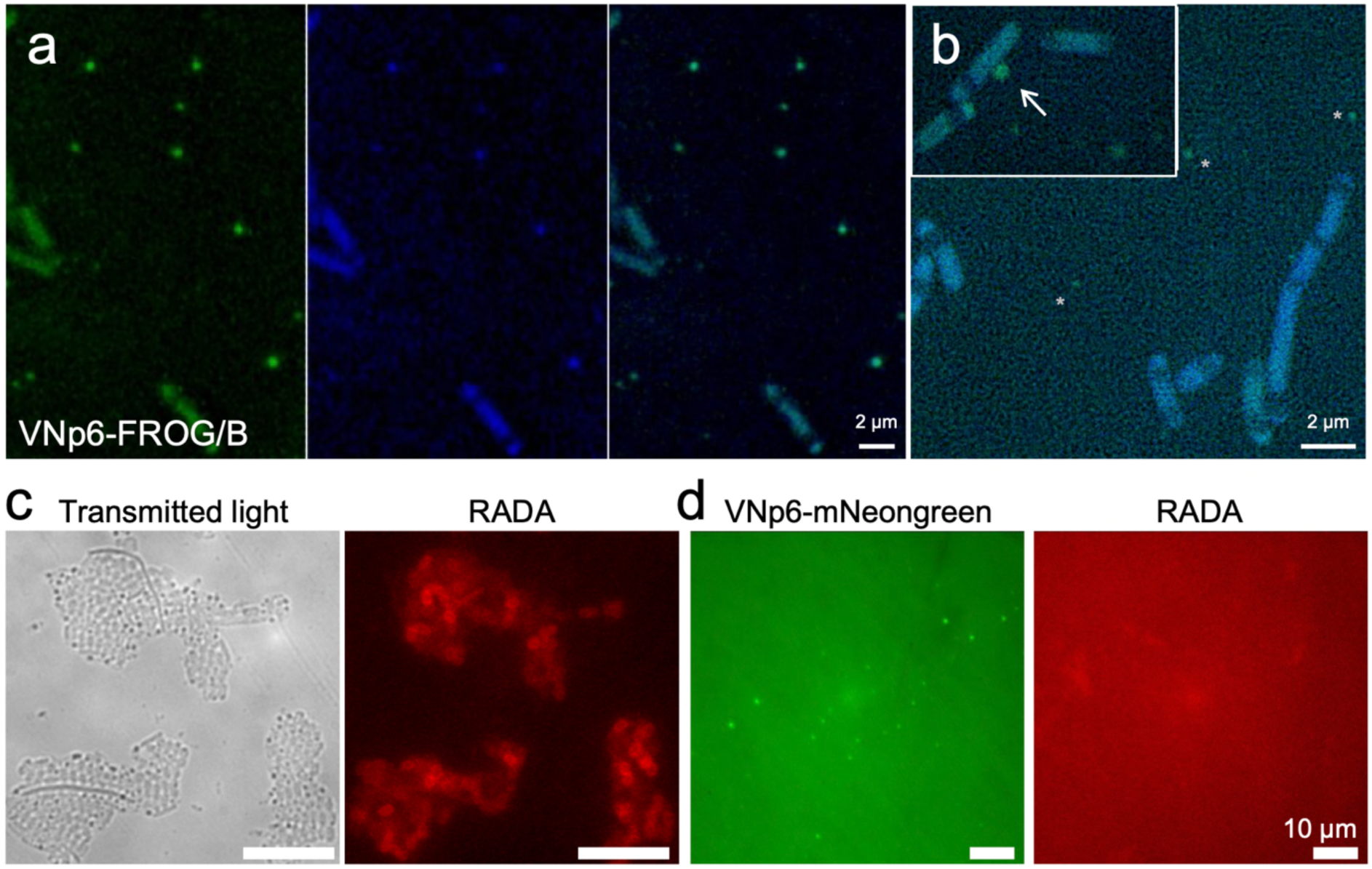
The environment within VNp induced extracellular vesicles. Widefield fluorescence (a) and Structured Illumination Microscopy (b) images of overnight cultures of *E. coli* cells expressing a fusion between VNp6 and the REDOX potential biosensor, FROG/B. These data illustrate the intra-vesicular environment is more oxidising (more green fluorescence) than the cytosol of the bacterial cells. Asterisks and arrow highlight vesicles in the SIM images. (c) While wide-field RADA staining (red) highlights the periplasmic localisation of peptidoglycan within *E. coli* cells, (d) the dye is not detected in isolated VNp-mNG (green) induced vesicles. (Scales in C: 10 µm)

We next compared yields of recombinant proteins produced within VNp induced vesicles and eOMVs. We monitored fluorescence of cultures of *E. coli* cells expressing VNp6 or ssDsbA tagged mCh2 within and without OmpX-mNG (Figure S5). The final cell density of each of the cultures were equivalent (not shown), indicating none of the constructs negatively impacted viability or growth of cells. In contrast cultures containing cells expressing VNp6-mCh2 had a maximal fluorescence at 589 nm six times higher than those expressing ssDsbA-mCh2. Interestingly while co-expression with OmpX-mNG had no significant impact on fluorescence dynamics in cells expressing ssDsbA-mCh2, co-expression with VNp6-mCh2 resulted in a higher overall maximal fluorescence, that remained stable after the culture reached stationary phase (Figure S5).

We examined how the overall culture fluorescence correlated to the intra-vesicular fusion proteins exported into the culture media by measuring the absorbance at the maximal extinction wavelength for the fluorescent proteins, mCh2 (589 nm) and mNG (506 nm) of media in which cells had been removed by centrifugation (Figure 5b). Average concentrations of vesicular fusion proteins within the culture media were subsequently calculated from a minimum of 3 independent biological repeats (Figure 5c). The VNp-fusion resulted in 298 ± 70 mg / litre yields of vesicle encapsulated mCh2 protein exported into the media, while the periplasmic targeting ssDsbA-fusion resulted in 42 ± 4 mg / litre yields. These data show that the VNp tag not only resulted in a higher yield of protein (Figure S5), but a significantly higher proportion of the resultant fusion protein was exported into vesicles compared to equivalent protein targeted to the *E. coli* periplasm. Interestingly, co-expression of OmpX-mNG (or unlabelled OmpX - not shown) with either VNp or ssDsbA fusions, resulted in a reproducible and significant increase in the extracellular yield of mCh2 (Figure 5c).

**Figure 5.**
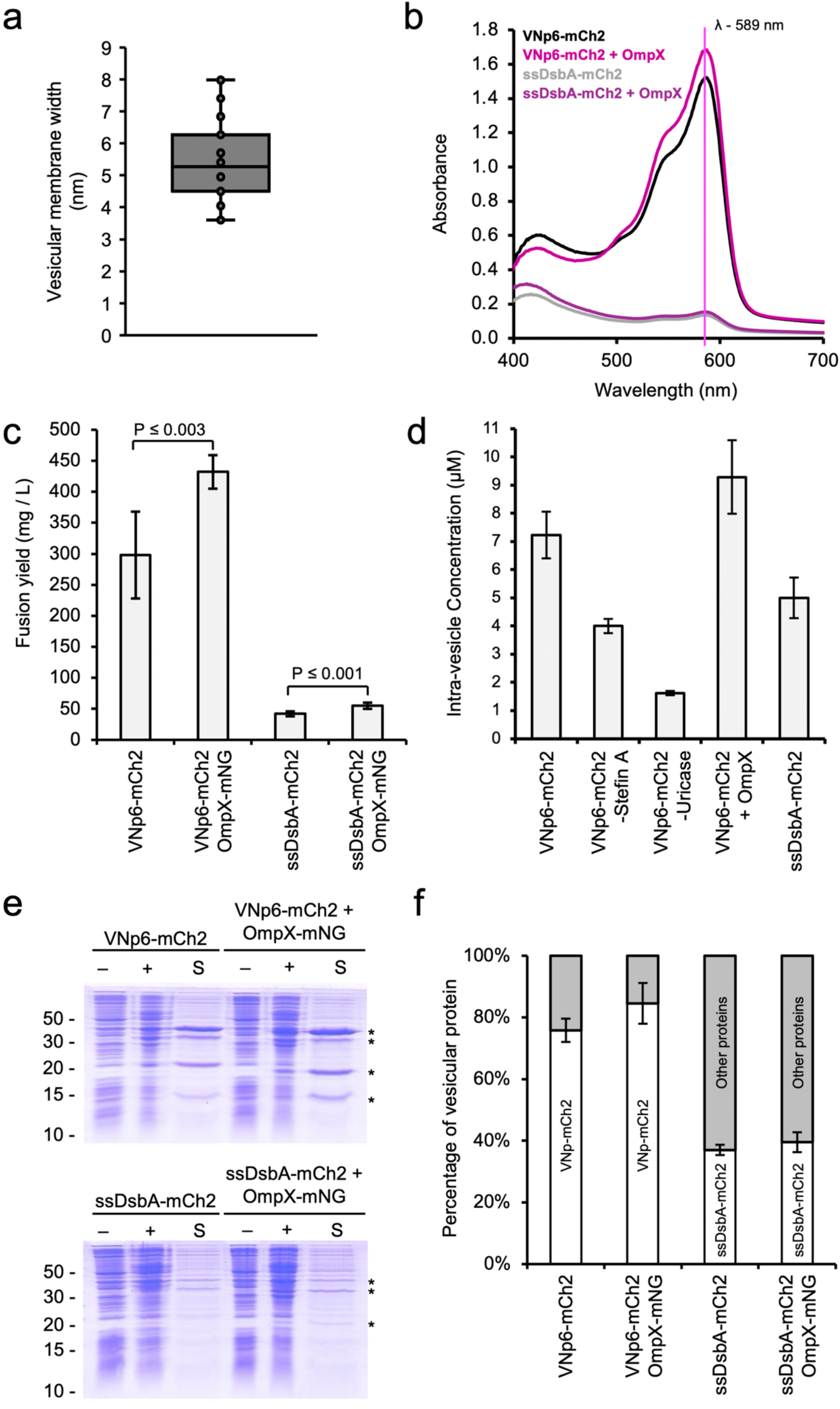
Characterisation of size and protein content of VNp vesicles and OMVs from *E. coli*. (a) Thickness of membrane of VNp6-mCh2 induced vesicles, determined from TEM thin section and Cryo EM data. (b) Typical absorbance spectra of media from cultures of cells expressing VNp6-Ch2 (black), VNp6-mCh2 & OmpX (pink), ssDsbA-mCh2 (grey) or ssDsbA & OmpX (purple). (c) Average yields calculated from spectra shown in (b), calculated from > 3 biological repeats. (d) Intra-vesicular concentration of mCh2 signal within vesicles isolated from cultures of *E. coli* expressing VNp6-mCh2, ssDsbA-mCh2, VNp6-mCh2 OmpX, VNp6-mCh2-StefinA or VNp6-mCh2-uricase, as determine by FLIM analysis. (e) Coomassie stained SDS-PAGE gels showing preinduction (-), overnight post-induction (+) cell pellet, and overnight supernatant (S) samples from cultures of *E. coli* expressing either VNp6-mCh2, ssDsbA-mCh2, VNp6-mCh2 OmpX-mNG, or ssDsbA-mCh2 OmpX-mNG. * - highlights bands that correlate with western blots of control mCh2 samples (Figure S5c). (f) Fraction (%) of total vesicular protein that is mCh2 (white) compared to other proteins (grey), averaged from 3 biological repeats of experiment show in (e).

Fluorescence Lifetime Imaging Microscopy (FLIM) was employed to determine whether these differences in protein yield was a result of differences between constructs in the quantities of fusion protein packaged into vesicles, or the number of vesicles produced. FLIM provides a robust method for determining concentrations of fluorescently labelled proteins within a sample (31, 32), taking into account the environment of the protein, such as protein-protein interactions and quenching. Steady-state imaging alone does not accurately give this information. Simply fluorescent proteins in highly viscous environment due to high protein concentration leads to reduction in lifetime. This method is not dependent on size of the object, as the concentration is calculated from the number of photons gathered at the defined pixel gate, and thus removes bias brought about by sample aggregation. Vesicular concentrations of VNp-fusions were determined using a previously described one photon excitation FLIM system (21, 30). Average photons / pixel gate was measured for ∼ 40 separate vesicles from 3 separate fields of view per sample, and average photon values were converted to fusion protein concentration using a calibration equation calculated using samples containing known concentrations of purified mCh2 protein. Using this method, we determined the intra-vesicle concentration for each fusion protein (Figure 5d, Table 1). Interestingly these values followed the same trends observed in yields (Figure 5c & d) increasing from ssDsbA (5 ± 0.72 µM), to VNp6-(7.2 ± 0.83 µM), to VNp6 & OmpX (9.3 ± 1.3 µM).

At the same time we examined the vesicular concentration of VNp-mCh2-StefinA (43.4 kDa) and VNp-mCh2-Uricase (66.8 kDa) fusion proteins, compared to VNp-mCh2 (30.9 kDa), to determine whether size of the fusion protein affected yield and intra-vesicle concentration of the VNp fusion. Stefin-A and Uricase were selected as both have previously been produced and exported at high yield with the VNp system (21, 51). A negative correlation was observed between the size of the fusion protein and both the intra-vesicular concentration (Figure 5d) as well as with the overall yield of each fusion protein from the culture (Figure S5b).

As well as establishing intra-vesicular concentration of the fusion proteins, we also used FLIM to determine viscosity within the different vesicles. Using a recently established technique (34), we examined the viscosity within vesicles generated from *E. coli* cells expressing either VNp6-mNG, ssDsbA-mNG, or co-expressing VNp6-mNG and OmpX (Table 1). mNG fusions were used, as this fluorescent protein is more sensitive to changes in viscosity than mCh2. Viscometry was the lowest in vesicles containing ssDsbA fusions (8.093 ± 0.011 centiPoise (cP)), increasing to 7.677 ± 0.020 cP for the VNp fusion (cP value increases inversely to viscosity), and was seen to be highest in vesicles containing VNp6-mNG and OmpX (5 cP). Thus, viscometry increases in relation to the intra-vesicular concentration of the fusion protein.

This finding is consistent with the VNp fusion making up a significant higher proportion of the overall vesicular content than the ssDsbA fusions within OMVs. To assess this further, we examined overall protein content of vesicles generated from the VNp and ssDsbA fusions, to determine differences in the relative purity of fusion between ssDsbA-fusion labelled OMVs and VNp induced vesicles. As a control, the identity of Coomassie stained mCh2 and mNG protein bands were confirmed by western blot analysis (Figure S5c). We reproducibly observed that VNp-fusions represent at least 80% of the total vesicle protein content (Figure 5e & f). In contrast the ssDsbA-fusion represented a significantly lower proportion of the overall protein content of eOMVs, which contained an abundance of endogenous *E. coli* proteins (Figure 5e). This data is consistent with natural blebbing due to increase in overall periplasmic content, rather than an induced nucleation and targeted vesicular delivery, observed with VNp-fusion encapsulated vesicle.

## Discussion

In this study we have compared recombinant Vesicle Nucleating peptide (VNp) induced vesicles (21) with endogenous Outer Membrane Vesicles (eOMV) from *E. coli*. The data indicates they are produced by distinct mechanisms, as VNp-fusions are absent from eOMVs, while periplasmic targeted proteins are not detected in VNp-induced vesicles. This exclusive segregation of molecules illustrates distinct mechanisms and is consistent with the observed differences in abundance and purity of the targeted fusion protein within the vesicles. Analysis of CryoEM and TEM images of VNp-mCh2 vesicles reveals these particles, are encapsulated within a single lipid bilayer. While VNp induced vesicles were occasionally observed with multiple membrane layers (in less than 1 in 20 vesicles) and therefore represents a small subset of vesicles containing both inner and outer membranes, potentially brought about as an artefact of EM sample preparation. The presence of Lipid-A and relative proportion of phosphatidylglycerol and cardiolipin (Figure 2), as well as the thickness of the lipid bilayer, strongly indicates that as for eOMVs, the VNp vesicle lipids originate from the outer *E. coli* membrane. This is consistent with the presence of outer membrane proteins within purified VNp induced vesicles, but no inner membrane proteins detected.(21)

While eOMV biogenesis is still not fully understood, genetic studies focussing on gene deletion strains have provided models to explain their formation. Strains lacking Lpp-peptidoglycan cross-links reduce vesiculation (52), whereas deletion of the outer membrane protein, OmpA, leads to increased OMV production. These effects are likely due to disruption between the OM and peptidoglycan (53, 54). It is probable OMV biogenesis requires a localised breakdown or disruption of the OM-peptidoglycan cross-links, otherwise it could only occur at regions where there are fewer crosslinks (i.e. at the cell poles). While eOMVs contain a high proportion of periplasmic proteins, only cytoplasmic and outer membrane proteins have been detected within VNp induced vesicles (21). These OM proteins and the presence of lipid-A provide compelling evidence that these vesicles form from the outer membrane. Full length recombinant Synucleins (from which the VNp tags are derived) has been reported to localise to the periplasm through the SRP-dependent pathway (55) when expressed recombinantly in *E. coli*. While no definitive mechanism for the translocation has been determined, with Sec and Tat pathways the predominant means by which proteins are transported to the periplasm (19), neither have been implicated here. Once the VNp fusion is translocated into the periplasm it must pass through the peptidoglycan by diffusion to reach the outer membrane. It has been postulated that globular proteins of up to 50 kDa are able to readily pass through pores in the stretched *E. coli* peptidoglycan layer (56), where they would be able to associate with the outer membrane and induce formation and incorporation into extracellular vesicles. This is consistent with our earlier published descriptions of the VNp system, that show larger proteins, and protein complexes tend to be incorporated into cytosolic vesicles (21, 23) (i.e. fail to pass across the peptidoglycan layer).

We postulate that, if the VNp peptide fusion is only able to access the inner membrane (i.e. is unable to pass the peptidoglycan layer), it induces curvature of this membrane into the cytoplasm rather than curvature of the OM into the media. This explains why sometimes both intracellular and extracellular vesicles are formed and that disulfide bond-containing proteins (e.g. VNp-mNG-Etanercept (84 kDa)) have been observed in both vesicle types showing that VNp fusions must be transported to the periplasm for this bond formation to occur. It is likely that molecular weight alone is an overly simplistic explanation for determining whether a VNp fusion will be exported or not. Other factors will have an impact, such as secondary/tertiary protein structure, overall shape and solubility. Consistent with this model, peptidoglycan would be absent from within the vesicles, as observed here (Figure 4d).

While Structured Illumination Microscopy timelapse imaging (Figure S4) does not provide the temporal and spatial resolution required to allow the capture of detailed membrane remodelling, it provides intriguing insights into the biomechanics of extracellular vesicle formation. To our knowledge this represents the first high-resolution timelapse description of vesicle formation from *E. coli* cells. Vesicles formed predominantly at the cell tips and site of cell division (Figure S4). Not only do the lipid composition, curvature, and dynamics of these membrane at these regions differ significantly from tubular regions of the cell, but the periplasm is also wider at these locations (49), allowing localised build-up of the VNp-fusion. Each of these factors are likely to be key factors in facilitating the rapid membrane reorganisation. Optimising conditions for the capture of vesicle formation highlighted key factors, other than minimizing the impact of phototoxicity (57, 58), required to promote vesicle formation. These include maximising cell health prior to and post induction; inducing at the optimal cell density (i.e. OD_600_ 0.8-1.0); as well as maintaining temperature, hydration, and maximising oxygenation throughout the experiment, key factors for maximising vesiculation in flask cultures. While the VNp fusion has some effect upon vesicle formation kinetics, with optimised conditions we observed the rate of vesicle production increases significantly 5 hours post induction.

We show that the small beta-barrel outer membrane protein, OmpX, localises to the surface of both VNp induced vesicles and eOMVs, consistent with the lipid bilayer originating from the bacterial outer membrane. Expression of additional OmpX or an OmpX-mNG fusion increases both VNp-fusion protein concentration within the vesicle as well as the overall yield within the culture. Our data suggests the additional OmpX does not impact the relative purity (Figure 5) of recombinant fusions within the VNp vesicles or eOMVs. Over-expression of OmpX does not bring about the formation of significantly larger vesicles (Table 1). Therefore, it is likely to be due to a combination of more efficient packaging within each vesicle, as well as overall increase in membrane stability.

In conclusion, the VNp-induced vesicles represent a distinct class of recombinant extracellular vesicles from eOMVs, that offer a simple and efficient route for producing high yields of relatively pure, concentrated, functional vesicle packaged recombinant protein, that can be applied to a wide range of protein production and biotechnological applications.

## Acknowledgements

The authors thank Gregory Mashanov for assistance with TIRF analysis of vesicles. This work was supported by the University of Kent and funding from the Biotechnology and Biological Sciences Research Council (BB/T008768/1 & BB/X007448/1). The authors acknowledge the use of Glacios 200 kV microscope, which received support from the Oncode Accelerator, a Dutch National Growth Fund project under grant number NGFOP2201. The Vesicle Nucleating peptide technology is described in the international patent filing *PCT/GB2022/053239*.

## Author contributions

B.R.S., T.A.E. and K.B. performed the experimental studies; M.L. performed TEM sample fixation, sectioning and imaging; A.E.B performed FLIM imaging; T.T.v.d.V. performed Cryo-EM sample fixation and imaging; L.J.C.J. supervised Cryo-EM analysis; S.W.B. undertook FLIM data analysis; L.W. supervised SIM microscopy; D.P.M. supervised research, sought funding and managed the overall project. All authors designed experiments. D.P.M. wrote main drafts of the manuscript, and all authors contributed to editing.

## Supplementary Figures

**Supplementary Figure 1.**
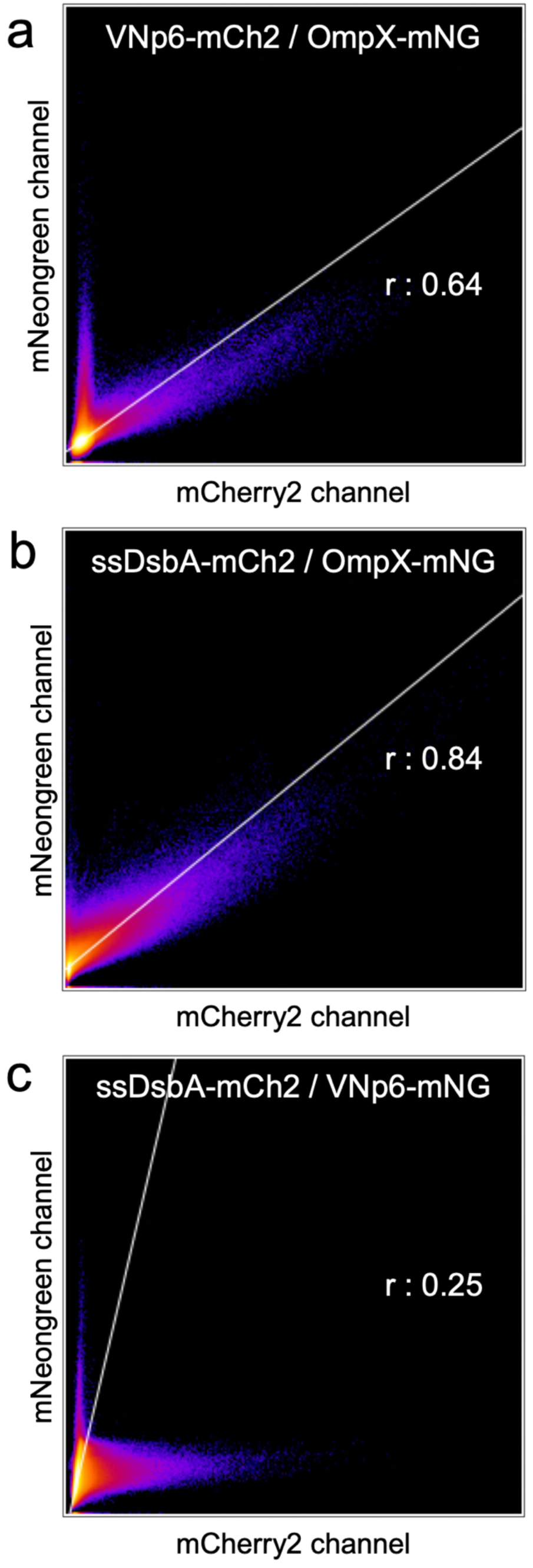
Colocalisation correlation analyses. Example colocalization correlation analysis graphs between mNG and mCh2 signals from Structure Illumination Microscopy images of vesicles isolated from BL21 DE3 *E. coli* expressing (a) VNp6-mCh2 and OmpX-mNG, (b) ssDsbA-mCh2 and OmpX-mNG, or (c) ssDsbA-mCh2 and VNp6-mNG, as shown in Figure 1. Perfect colocalization between the green (y-axes) and red (x-axes) channels would present as a distribution that follows a line of equality, with a calculated Pearson correlation value (r) of 1.

**Supplementary Figure 2.**
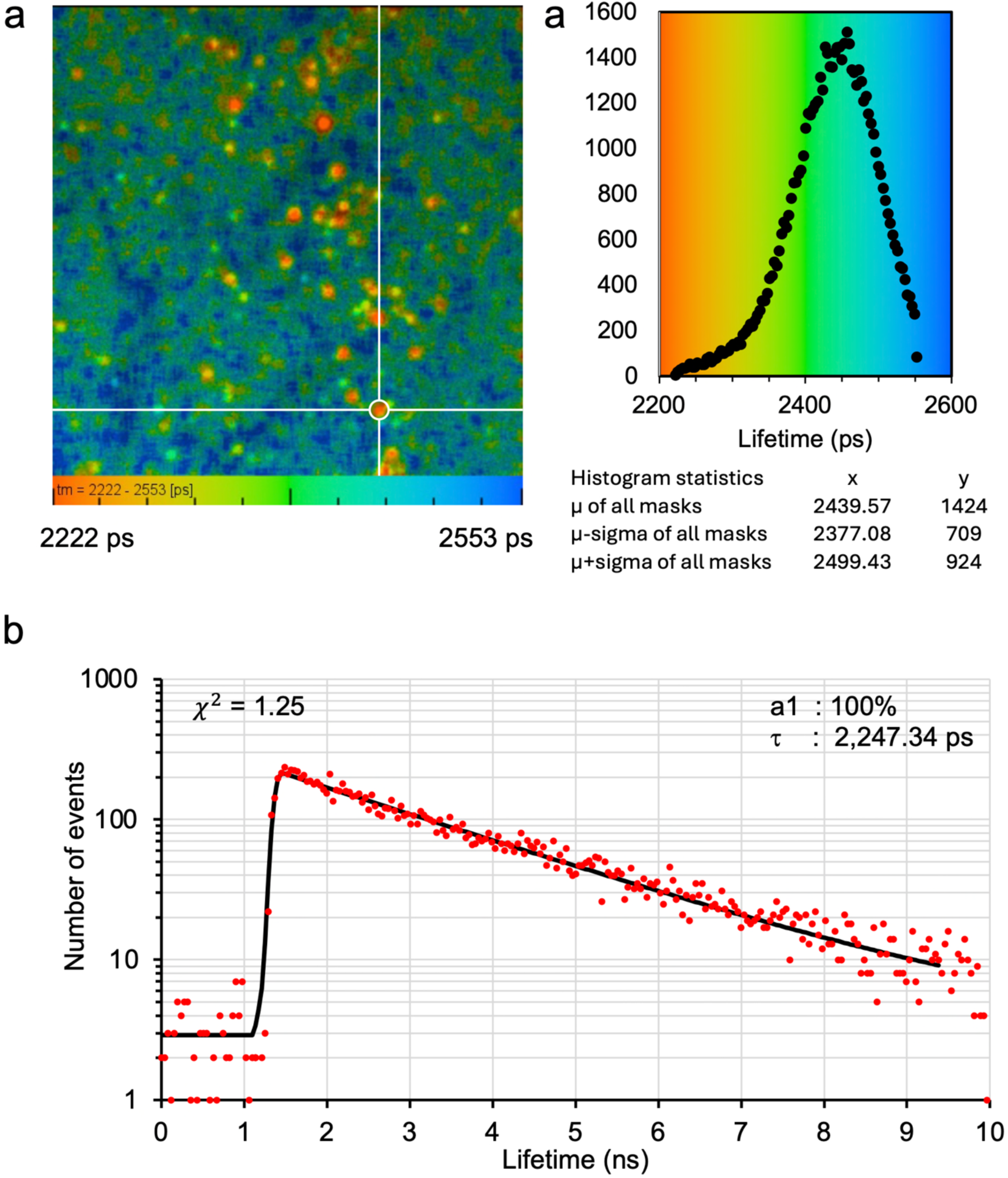
Example calculated lifetime of FLIM acquisition of vesicles isolated from *E. coli* BL21 DE3 cells expressing VNp6-mNG. (a) Analysed colour-coded FLIM map of VNp6-mNG containing vesicles. (b) Lifetime distribution histogram of total image from A. (c) Single exponential decay fit (pre-exponential factor a-100%), using 3x binning, of a pixel within highlighted circle within (a). Calculated lifetime (τ) of selected pixel is 2,247 ps. χ^2^ value of 1.25 indicative of a perfect single decay fit (black) to the data (red).

**Supplementary Figure 3.**
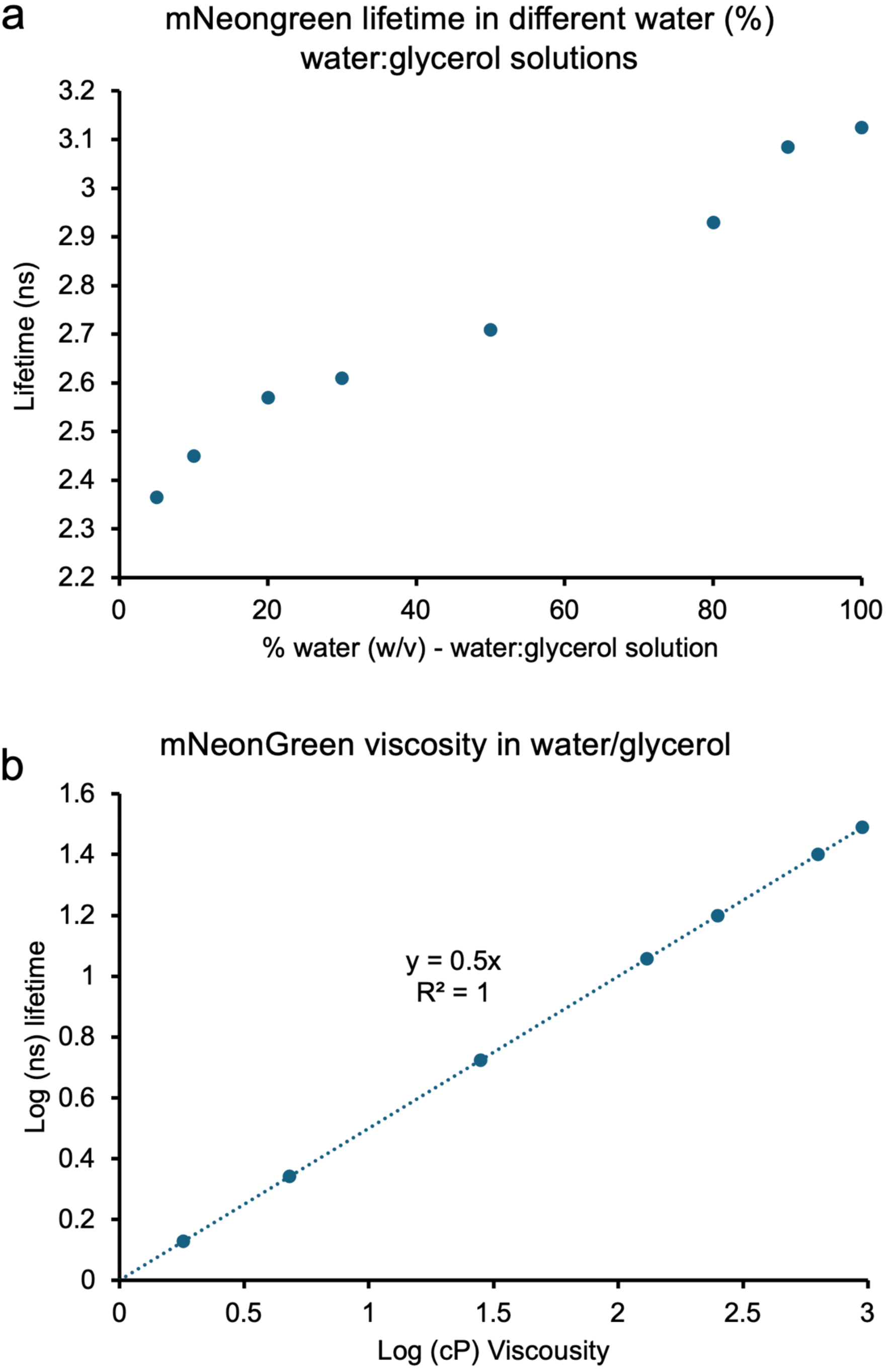
Calibration of viscosity dependent changes in mNG fluorescence lifetime. An mNG concentration calibration equation generated from photon measurements averaged from 5 separate FLIM acquisitions of equivalent samples of purified fluorescent dissolved in different water: glycerol ratio solutions.

**Supplementary Figure 4.**
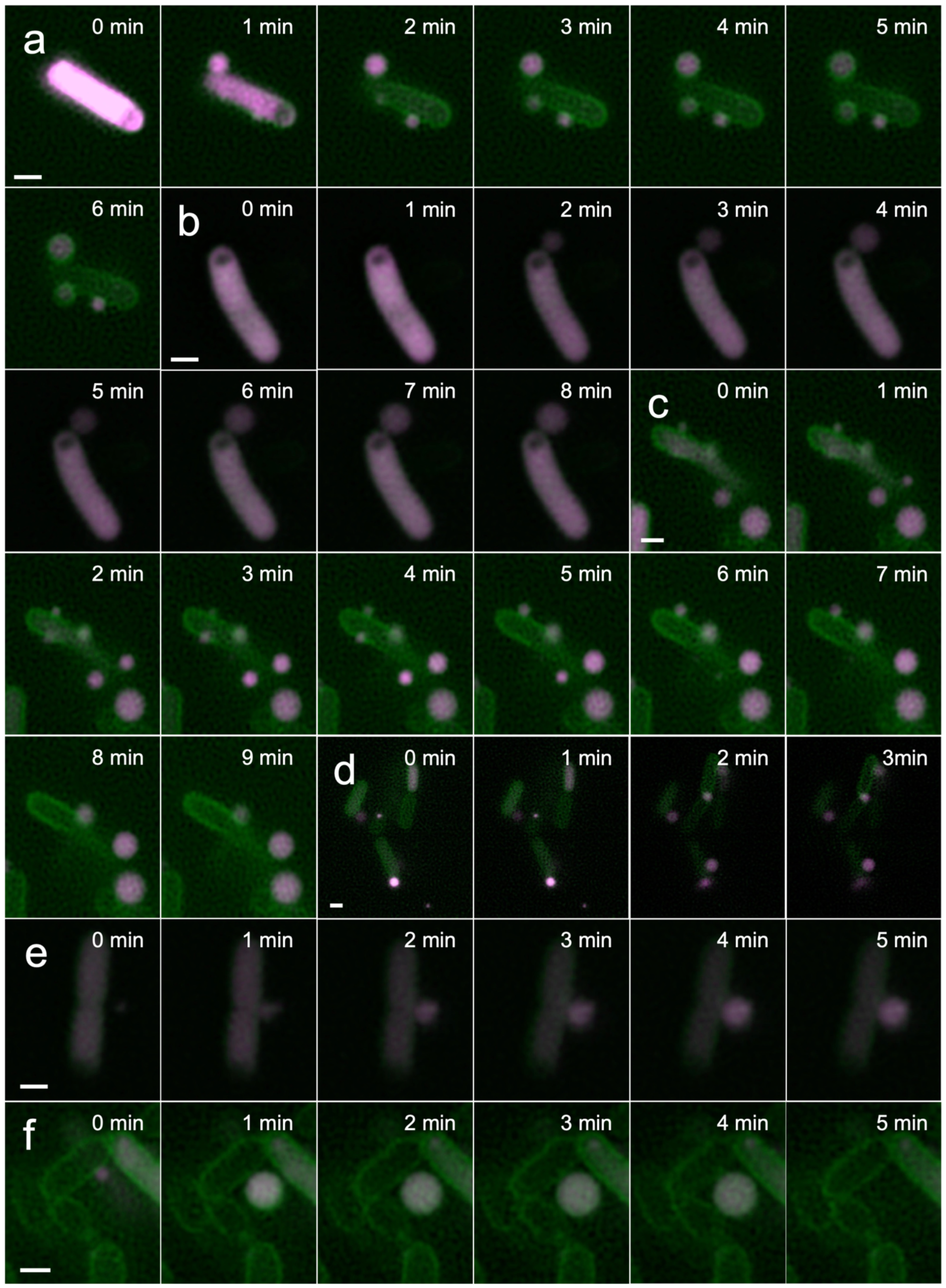
Frames from Structured Illumination Microscopy Timelapse videos of VNp-vesicle formation. 1 minute time frames from Structured Illumination Microscopy timelapse videos of live *E. coli* co-expressing VNp6-mCh2 (magenta) and the outer membrane marker OmpX-mNG (green), from 6 individual cells (a - f). Scale bars: 1 µm.

**Supplementary Figure 5.**
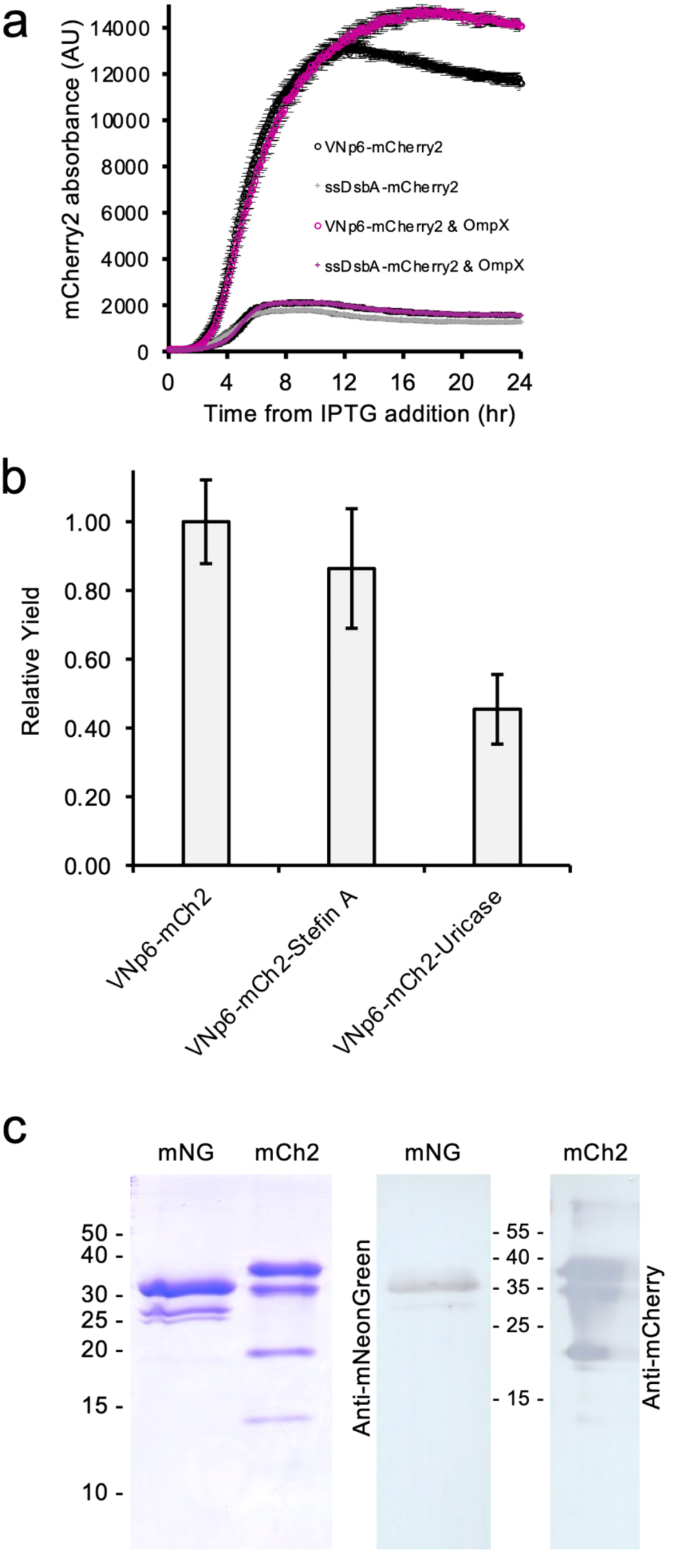
Yield of vesicular packaged recombinant fluorescent protein tagged fusions. (a) Fluorescence from overnight cultures of BL21 DE3 cells expressing VNp6-mCh2 (black circles), ssDsbA-mCh2 (grey crosses), VNp6-mCh2 OmpX-mNG (magenta circles), or ssDsbA-mCh2 OmpX-mNG (magenta crosses). Averages values as presented were calculated from 4 separate biological repeats. (b) Average yields of exported VNp-mCh2, VNp-mCh2-StefinA and VNp-mCh2-Uricase, each calculated from > 3 independent biological repeats. (c) Coomassie stained SDS-PAGE (left) and western blot (right) analysis of Ni^2+^ affinity purified mNG and mCh2 proteins. Commercial polyclonal anti-mNG and mCh2 primary antibodies (see materials and methods) were used.

